# Complete Genomic Characterization of Global Pathogens, Respiratory Syncytial Virus (RSV), and Human Norovirus (HuNoV) Using Probe-based Capture Enrichment

**DOI:** 10.1101/2024.09.16.613242

**Authors:** Sravya V Bhamidipati, Anil Surathu, Hsu Chao, Daniel P Agustinho, Qin Xiang, Kavya Kottapalli, Abirami Santhanam, Zeineen Momin, Kimberly Walker, Vipin K Menon, George Weissenberger, Nathanael Emerick, Faria Mahjabeen, Qingchang Meng, Jianhong Hu, Richard Sucgang, David Henke, Fritz J Sedlazeck, Ziad Khan, Ginger A Metcalf, Vasanthi Avadhanula, Pedro A Piedra, Sasirekha Ramani, Robert L Atmar, Mary K Estes, Joseph F Petrosino, Richard A Gibbs, Donna M Muzny, Sara Javornik Cregeen, Harsha Doddapaneni

## Abstract

Respiratory syncytial virus (RSV) is the leading cause of lower respiratory tract infections in children worldwide, while human noroviruses (HuNoV) are a leading cause of epidemic and sporadic acute gastroenteritis. Generating full-length genome sequences for these viruses is crucial for understanding viral diversity and tracking emerging variants. However, obtaining high-quality sequencing data is often challenging due to viral strain variability, quality, and low titers. Here, we present a set of comprehensive oligonucleotide probe sets designed from 1,570 RSV and 1,376 HuNoV isolate sequences in GenBank. Using these probe sets and a capture enrichment sequencing workflow, 85 RSV positive nasal swab samples and 55 (49 stool and six human intestinal enteroids) HuNoV positive samples encompassing major subtypes and genotypes were characterized. The Ct values of these samples ranged from 17.0-29.9 for RSV, and from 20.2-34.8 for HuNoV, with some HuNoV having below the detection limit. The mean percentage of post-processing reads mapped to viral genomes was 85.1% for RSV and 40.8% for HuNoV post-capture, compared to 0.08% and 1.15% in pre-capture libraries, respectively. Full-length genomes were>99% complete in all RSV positive samples and >96% complete in 47/55 HuNoV positive samples—a significant improvement over genome recovery from pre-capture libraries. RSV transcriptome (subgenomic mRNAs) sequences were also characterized from this data. Probe-based capture enrichment offers a comprehensive approach for RSV and HuNoV genome sequencing and monitoring emerging variants.

**IMPORTANCE:** Respiratory syncytial virus (RSV) and human noroviruses (HuNoV) are NIAID category C and category B priority pathogens, respectively, that inflict significant health consequences on children, adults, immunocompromised patients, and the elderly. Due to the high strain diversity of RSV and HuNoV genomes, obtaining complete genomes to monitor viral evolution and pathogenesis is challenging. In this paper, we present the design, optimization, and benchmarking of a comprehensive oligonucleotide target capture method for these pathogens. All 85 RSV samples and 49/55 HuNoV samples were patient-derived with six human intestinal enteroids. The methodology described here results has a higher success rate in obtaining full-length RSV and HuNoV genomes, enhancing the efficiency of studying these viruses and mutations directly from patient-derived samples.

## INTRODUCTION

Respiratory syncytial virus (RSV) and human norovirus (HuNoV) are clinically significant pathogens due to the considerable burden of disease they impose globally(1, 2). RSV is the leading cause of severe respiratory illness and mortality especially in infants and young children, and a major cause of illness in the elderly(3). HuNoV is the most common cause of acute gastroenteritis globally(4). While all viruses warrant attention in virology and public health, the high prevalence and broad impact of RSV and HuNoV infections underline their particular importance.

RSV and HuNoV are RNA viruses, with distinctive genome structures and characteristics that define their respective families(5,6). RSV belongs to *Pneumoviridae* family and *Orthopneumovirus* genus and carries a single-stranded, negative-sense, non-segmented RNA genome. The RSV genome consists of approximately 15,200 bp containing 10 genes encoding 11 proteins. Each gene encodes for a separate mRNA except M2, which contains two overlapping open reading frames (ORFs) (5). HuNoV is a positive-sense, single-stranded RNA virus that belongs to the *Caliciviridae* family. The genome is between 7,500 to 7,700 bp in length and is divided into three overlapping ORFs(7) ORF1 encodes a large polyprotein cleaved into six non-structural proteins, while ORF2 and ORF3 encode the major (VP1) and minor (VP2) capsid proteins respectively. The HuNoV genome is covalently linked at the 5’ end to a small viral protein (VPg), which is instrumental for the initiation of protein synthesis(6, 8, 9), and is polyadenylated at the 3′-end.

RSV and HuNoV are known for their substantial strain diversity(3, 9) and are divided into numerous genotypes, each bearing unique genetic sequences. RSV is divided into two major subtypes: RSV-A and RSV-B, based on major antigenic differences in the G glycoprotein and reactivity to monoclonal antibodies(10, 11). These groups are further classified into genotypes based on the nucleotide sequence of the second hypervariable region of the C-terminal end of the G gene. The number of RSV genotypes keeps evolving, with 24 lineages within RSV-A and 16 within RSV-B identified thus far(12, 13). However, there is no consensus on the classification for assigning genotypes or their nomenclature. The most recent genotypes circulating worldwide are RSV/A/Ontario (ON) and RSV/B/Buenos Aires (BA), with a unique 72 and 60 nucleotide duplication in the distal third of the G gene, respectively. Based on phylogenetic analysis of major capsid protein VP1 amino acid sequences, noroviruses are divided into ten genogroups (GI-GX), of which human infections are caused by viruses in GI, GII, GIV, GVIII, and GIX genogroups. Each genogroup is divided into genotypes and some genotypes are further divided into variants. The prototype HuNoV is the GI.1 Norwalk virus. GII.4 viruses are responsible for a majority of the HuNoV outbreaks worldwide(8, 13). Although other genotypes such as GII.17 have emerged as the leading cause of gastroenteritis in some countries in some years(14). Therefore, obtaining full-length genomes to facilitate accurate characterization of RSV and HuNoV genotypes is important to monitor their epidemiology.

There are several demonstrated approaches to obtain genomic sequences from viruses(15). RSV sequencing has been reported using NGS methods such as overlapping amplicon-based and targeted metagenomic sequencing(16–19). For HuNoV, amplicon-based sequencing(20), capture probe-based enrichment(21, 22), PolyA+ enrichment (23) and long read sequencing(24) have been described. Each of these methods has its caveats, and obtaining full-length genomes from these viruses has been challenging due to the sequence heterogeneity among different genotypes and low viral titers in some samples. Furthermore, the current commercial options such as the Twist Comprehensive Viral Research Panel, for capture-based enrichment are designed to enrich and detect a broad range of viruses rather than targeting RSV and HuNoV viruses and all their known genotypes for complete genome sequencing(25). This study aims to provide comprehensive probe sets for these two important viral pathogens and a single workflow that can be used to recover full-length genomes and facilitate accurate genotyping of both viruses. Furthermore, the generated sequence data has been demonstrated for the first time to study the RSV genome ORF expression patterns.

## RESULTS

We utilized capture probes and a streamlined target enrichment workflow for sequencing and analysis of RSV and HuNoV genomes (Fig.1). To demonstrate the utility of the capture enrichment methodology, sequencing data from pre-and post-capture libraries of both RSV and HuNoV were analyzed for efficiency of genome recovery and accuracy of genotyping. Samples used in this study were all RSV or HuNoV positive and their subtypes/genotypes were previously determined using qPCR assays as detailed in the methods. For RSV, 85 post-capture libraries and 24/85 pre-capture libraries belonging to RSV-A and RSV-B subtypes were sequenced (Table 1). For HuNoV, 55 post- and pre-capture libraries were sequenced. These 55 HuNoV represent GI.1, GII.4, and other GII genotypes (GII.3, GII.6, and GII.17) (Table 1).

**Fig. 1.**
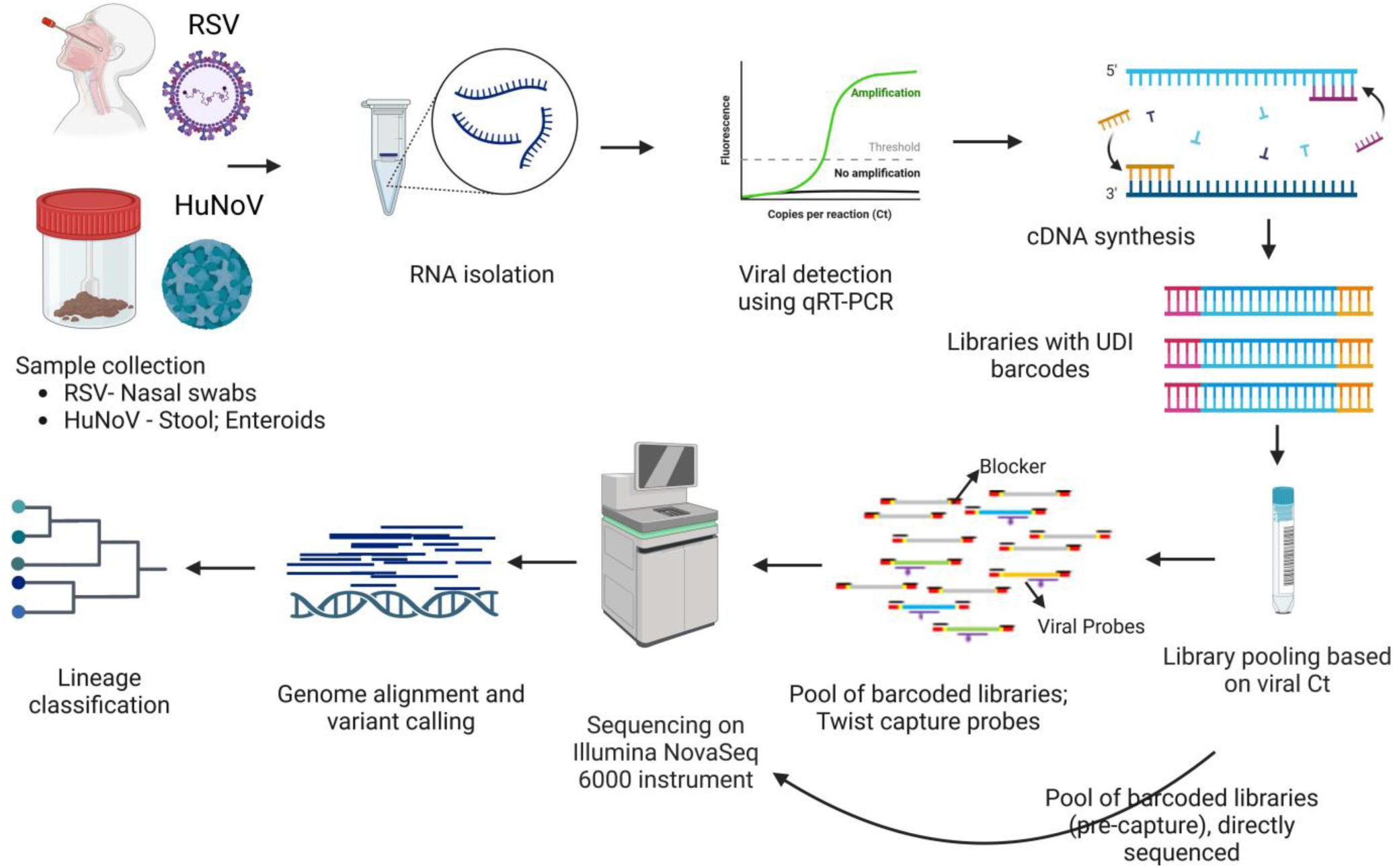
Schematic workflow. Presented in the workflow are the different steps involved in the RSV and HuNoV capture and sequencing methodology. First row—RNA was isolated from mid-turbinate nasal swab samples (RSV) and from stool samples or infected human intestinal enteroids (HuNoV) followed by Real-Time RT-PCR to detect these viruses. Positive samples were quantified, and RNA was converted to cDNA. Second row–The cDNA was used to generate Illumina libraries with molecular barcodes and these libraries were pooled based on the Ct. values. Capture enrichment was performed with either RSV or HuNoV probe set, and enriched libraries were then sequenced on the Illumina NovaSeq 6000 instrument to generate 2×150 bp length reads. Pre-captured libraries were also sequenced followed by downstream genome reconstruction, variant, and lineage analyses.

**Table 1.**
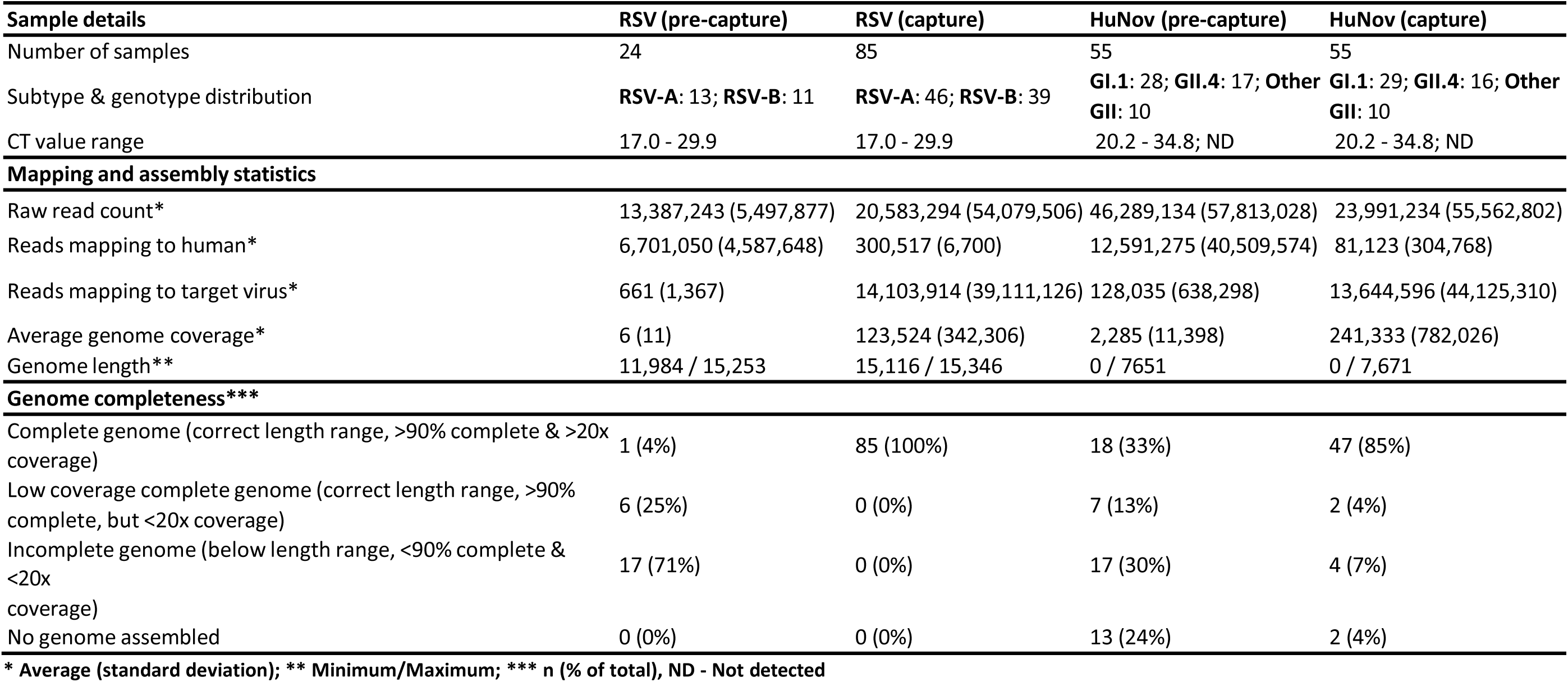
Sample composition, mapping, and genome assembly statistics.

**Table 2.** The number of isolates used and the final capture probe design details.

|  | RSV | HuNoV |
| --- | --- | --- |
| **Strains** | **A&B** | **GI, GII, GIV** |
| **# of Seq (Isolates)** | **1,570** | **1,376** |
| **Bases Targeted** | **23.77 Mb** | **9.6 Mb** |
| **% Bases Covered** | **99.79** | **99.68** |
| **Unique Probes (80 bp)** | **87,025** | **39,300** |

### Sequencing results and capture enrichment efficiency

The sequences were trimmed to remove low-quality regions, and the resulting non-human reads were analyzed using the VirMAP pipeline (24). A summary of the mapping and assembly statistics can be found in Table 1 and Table S1. Overall, most post-processing reads in the post-capture libraries mapped to their respective target virus; this proportion was significantly lower in pre-capture libraries (Fig. 2).

**Fig. 2.**
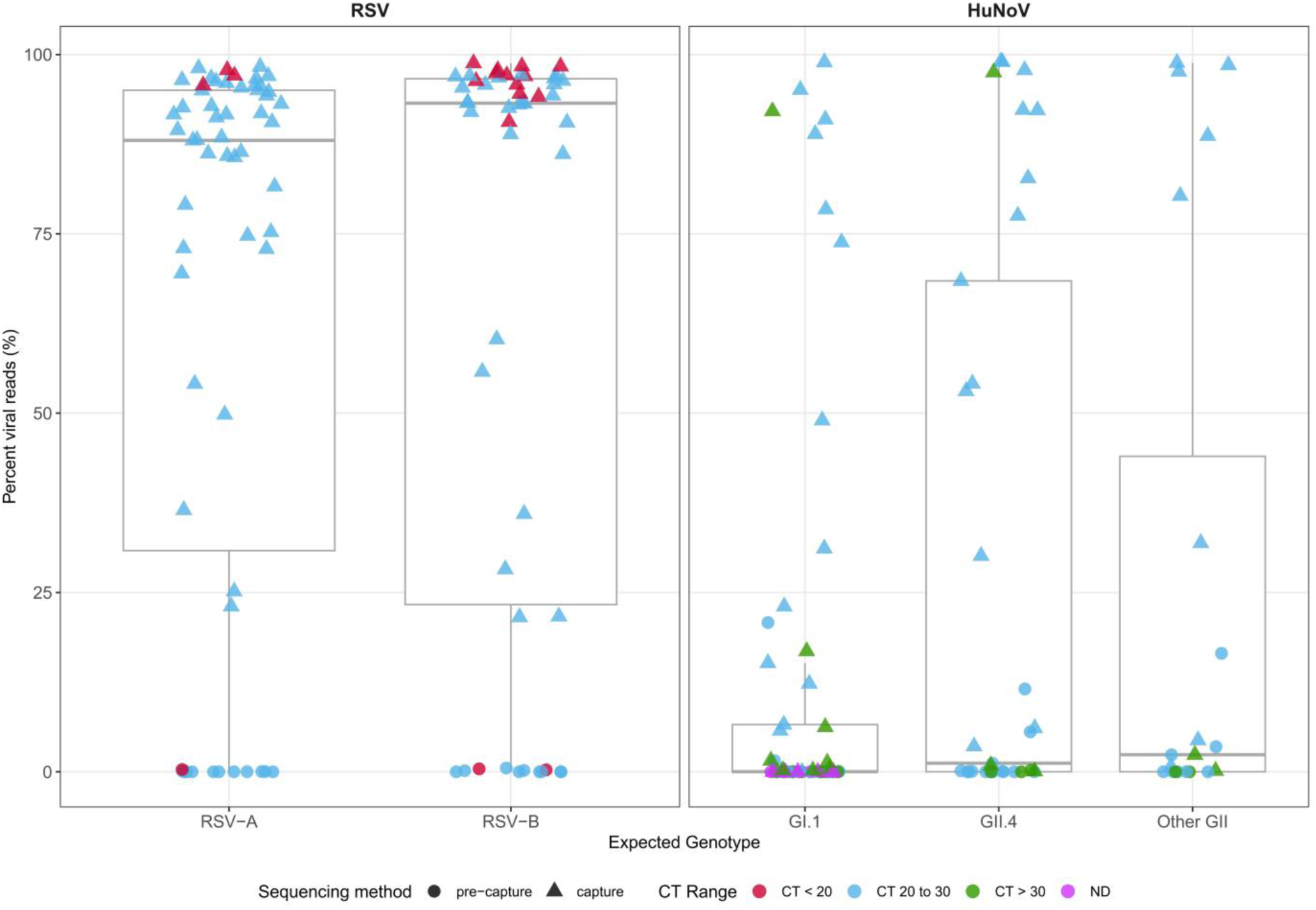
Viral read recovery efficiency. Percent of trimmed, non-human sequence reads (post-processing) that mapped to the target viral genome in pre-capture (circles) and post-capture (triangles) libraries. CT value range of samples: ‘CT <20’ (red), ‘CT 20 to 30’ (light blue), ‘CT > 30’ (green) & ND (not detected) (pink). A: Viral reads mapping to RSV genomes, split by two subtypes. B: Viral reads mapping to HuNoV genomes, split by genotypes (GI.1, GII.4, Other GII).

A total of 1.74 billion raw reads were generated from 85 RSV post-capture libraries with an average of 20.58 million (SD = 54 million) total raw, 300,000 (SD = 6,7000) host-mapped, and 14 million (SD = 39.1 million) viral genome mapped reads. (Table 1). The mean percentage of post-processing reads mapped to the RSV genome was 85.1%. This pattern was similar between RSV-A and RSV-B subtypes (Fig. 2). To assess the enrichment efficiency of post-capture libraries compared to pre-capture libraries, a subset of 24 pre-capture libraries were randomly selected and sequenced they generated a total of 0.32 billion raw reads with an average of 13.3 million (SD= 5.4 million) total raw, 6.7 million (SD= 4.5 million) host-mapped, and 661 (SD= 1,3000) RSV mapped reads (Table 1). The mean percentage of post-processing reads mapped to the RSV genome in the pre-capture libraries was 0.08% (Fig. 2).

The 55 HuNoV post-capture libraries generated a total of 1.31 billion raw reads with an average of 23.9 million (SD = 55.5 million) total raw, 81123 (SD = 304,000) host mapped, and 13.6 million (SD = 44.1 million) HuNoV mapped reads (Table 1). To assess the capture efficiency 55 pre-capture libraries were sequenced. They generated a total of 2.54 billion raw with an average of 46.2 million (SD= 57.8 million) total raw, 12.5 million (SD = 40.5 million) host mapped, and 128,000 (SD = 638,000) HuNoV mapped reads (Table 1). The mean percentage of post-processing reads mapped to HuNoV genomes was 40.8% in post-capture libraries and 1.15% in the pre-capture libraries. The percentage of reads that mapped to the HuNoV genomes varied among the genotypes as shown in (Fig. 2). Detailed statistics for RSV and HuNoV genomes can be found in Table S1.

### The comprehensiveness of genome recovery and genotyping

To evaluate the capability of the capture methodology to assemble full-length genomes, the VirMAP pipeline was used to reconstruct RSV and HuNoV genomes. The VirMAP summary statistics are shown in Fig. 3 and Table 1. Genome recovery success using the capture probe sets was evaluated, by classifying the genome reconstruction as **‘complete’** (within expected length range, >90% completeness & >20x coverage), **‘complete with low coverage’** (within expected length range, >90% completeness & <20x coverage) or **‘incomplete’** (below expected length range, <90% completeness & <20x coverage).

**Fig. 3.**
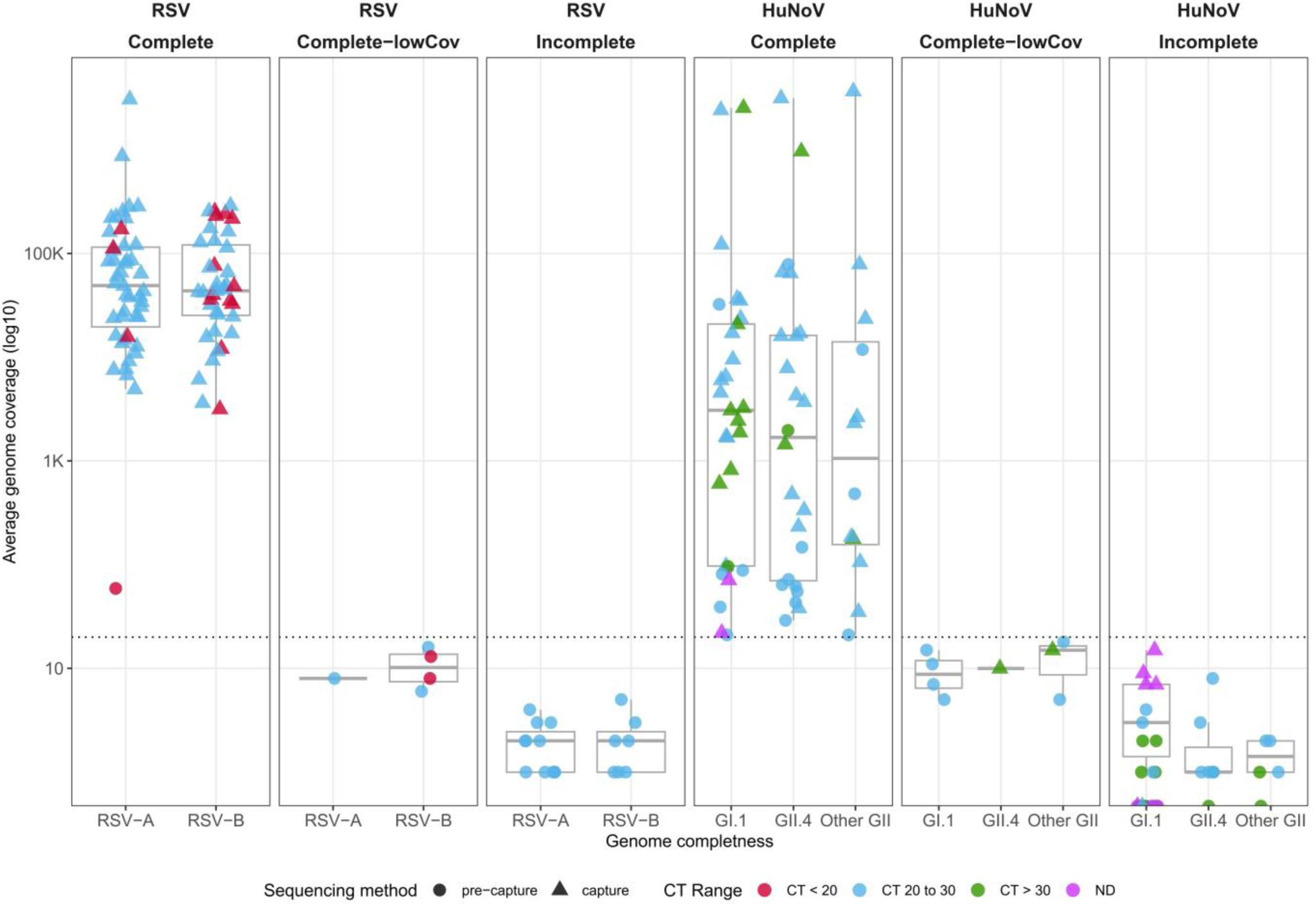
Average genome coverage obtained in post-capture (triangles) and pre-capture (circles) samples. Genome reconstruction was classified as follows: ‘complete’ (within expected length range, >90% completeness & >20x coverage), ‘complete with low coverage’ (within expected length range, >90% completeness & <20x coverage), or ‘incomplete’ (below expected length range, <90% completeness & <20x coverage). CT value range of samples: ‘CT <20’ (red), ‘CT 20 to 30’ (light blue), ‘CT > 30’ (green) & ‘ND’ (pink). A: RSV samples split by RSV-A or RSV-B genotype. B: HuNoV samples split by five genotypes (GI.1, GII.4, Other GII).

Complete genomes were successfully reconstructed for all 85 post-capture RSV libraries. In the 24 pre-capture libraries, there was one complete genome, six complete with low coverage, and 17 incomplete genomes (Fig. 3). The assembled genome length for the post-capture libraries was between 15,116 and 15,346 bp, and between 11,948 and 15,253 bp in pre-capture libraries (Table 1). The average coverage ranged from 3,153x to 3.05 million x with a mean of 123,000 (SD= 342,000) in post-capture. In 24 pre-capture libraries, it ranged from 1x to 59x, with a mean of 6x (SD =11) (Fig. 3 and Table S1). The 85 RSV post-capture genomes had a completeness of 99-100%, allowing the assignment of subtype as RSV-A or RSV-B (Table S1).

Of the 55 HuNoV post-capture libraries, 47 yielded complete genomes. Of the remaining eight samples, two samples resulted in low coverage complete genomes; four had incomplete genomes, and in the remaining two samples, genome assembly failed (Fig. 3 and Table 1). Sample p1540-BCM18-4 with a Ct value of 30.4 produced a low coverage (10x) complete genome and sequencing of the pre-capture library recovered an incomplete (12.9%) genome at only 1x coverage. Similarly, a low coverage complete genome (90% and 15x) was recovered from sample p1540-BCM18-5-AP however this sample had a high Ct value of 34.4. The four samples with incomplete genomes had Ct values ranging from 34.5 to Ct below the detection limit. The remaining two samples that failed to produce genome assemblies had Ct values of 28.3 and below the detection limit, respectively, and both underperformed in the pre-capture libraries, pointing to sample-related issues.

Of the 55 HuNoV pre-capture, 18 samples yielded complete genomes. There were 7 samples with complete low coverage, 17 with incomplete genomes, and 13 samples for which the genome assembly failed (Fig. 3 and Table 1).

The assembled genome lengths of the HuNoV post-capture libraries were between 0 and 7,671 bp and for pre-capture libraries between 0 and 7,651 bp (Table 1). The genome coverage ranged from 0x to 3.64 million x, with a mean of 241,000 x (SD = 782,000) in the post-capture libraries. The pre-capture libraries yielded a genome coverage range between 0 – 78,000x, with a mean of 2,284x (SD = 113,000) (Table 1 and Table S1).

Complete HuNoV genome reconstructions were genotyped via the CDC-developed Human Calicivirus Typing Tool (https://calicivirustypingtool.cdc.gov/bctyping.html). Of the 47 samples with complete genomes, 22 belonged to GI.1, 15 belonged to GII.4 and the remaining 10 belonged to other GII genotypes. (Fig. 3 and Table 1). In both RSV and HuNoV data sets, there was agreement in subtype or genotype assignment between the complete post-capture and pre-capture genomes.

To assess the ability of this probe-based capture enrichment method to enhance viral genome coverage depth, we realigned reads to either a reference genome (RSV) or individual sample-assembled genomes (HuNoV) and calculated the percentage of bases in the genome that are covered at a minimum of 20x in both post- and pre-capture libraries. Through this analysis, three HuNoV samples that met the first genome completeness criteria showed a relatively low breadth of 20x coverage. (Fig 4). To rule out any process-related issues or problems with the capture probe itself, fresh pre- and post-capture libraries were sequenced for these three samples (p1540-723-100595-AP, p1540-TCH-17-78-AP, and p1540-BCM18-5-AP). The results were the same as the first time, indicating that the problem is sample-related.

**Fig. 4.**
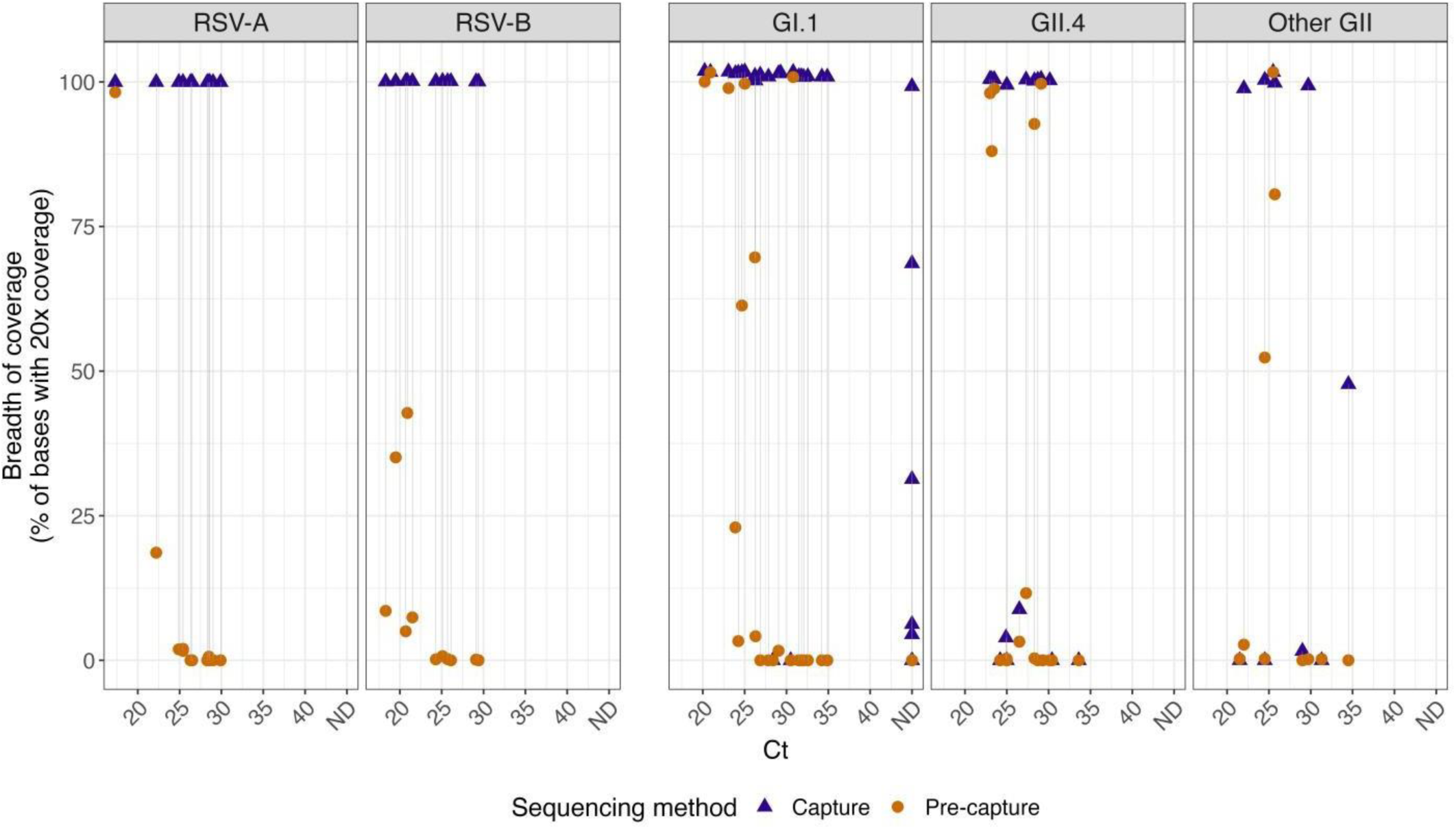
The breadth of coverage for a minimal 20x coverage was calculated from the post-capture and pre-capture RSV and HuNoV libraries. Sample pairs (i.e. the same sample processed with or without capture) are shown connected by a line. Samples that could not be detected by PCR were represented with ND (not detected). The left panel represents RSV and the right panel HuNoV subgroups.

### RSV ORF expression

To identify and quantitate sub-genomic mRNAs, the sequenced RSV reads were aligned to RSV-A or RSV-B reference genomes. The RSV genome has a total of 11 ORFs and the ORF read coverage for genotypes RSV-A and RSV-B are presented as normalized read pair counts (FPKM-reads per kilobase million) (Fig. 5).

**Fig. 5.**
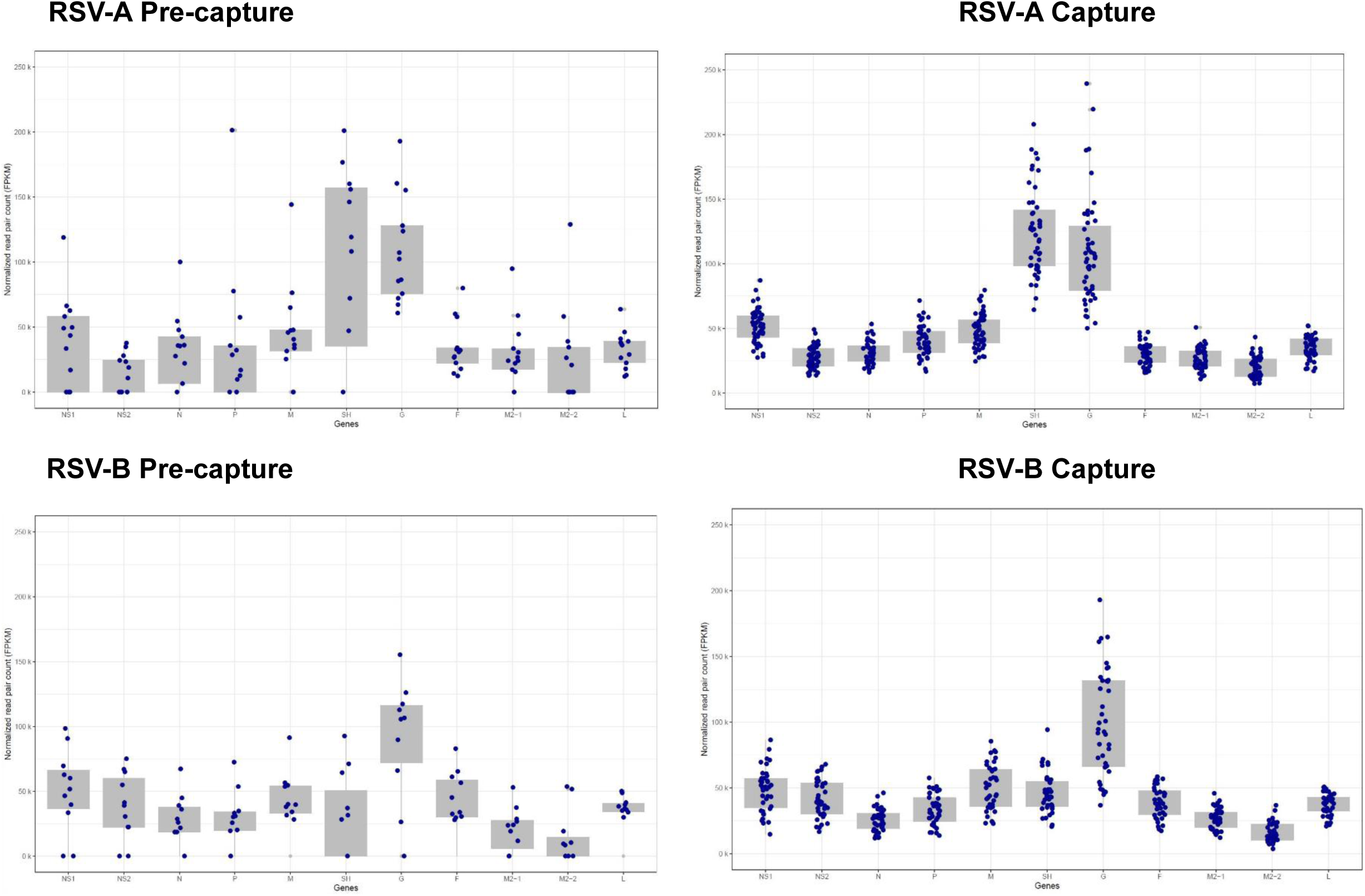
ORF expression levels in RSV-A (top panels) and RSV-B; (lower panels) pre-capture (left panels) and post-capture (right panels) samples.

A total of 46 samples were infected with RSV-A subtype. All 11 ORFs were quantified in post-capture libraries (Fig. 5). ORFs SH and G had the highest expression with an average of 124,303 and 109,011 FPKM respectively (Table S2). ORF M2-2 & M2-1, on the other hand, had the lowest expression with 19,890 and 26,690 FPKM respectively.

In comparison, 13 pre-capture libraries belonging to the RSV-A genotype, ORFs SH and G showed the highest expression, with an average of 139,449 and 109,086 FPKM respectively. The lowest expression was seen in ORFs NS2 and M2-2, with an average of 13,659 and 23,684 FPKM, respectively. Incomplete expression of ORFs was recorded in 9 pre-capture libraries, likely due to low read coverage. Notably, NS2 and M2-2 were not detectable in 7 and 6 of the pre-capture libraries, respectively (Table S2).

The remaining 39 samples were infected with the RSV-B subtype, all 11 ORFs were expressed in post-capture. ORFs G and M had the highest average FPKM values of 98,558 and 49,966, respectively, and ORFs M2-2 and N had the lowest values of 16,173 and 25,708 FPKM, respectively (Fig. 5).

In the 11 pre-capture libraries, the expression level was highest in ORFs G and NS1 with average values of 106,693 and 50,260 FPKM, and the lowest values of 13,829 and 20,336 were in ORFs M2-2 and M2-1 respectively. In 6 pre-capture libraries, incomplete expression of ORFs occurred. Expression was not detected in 5 libraries for ORFs M2-2 and M2-1, while SH ORF expression was not detected in 4 libraries (Table S2).

## DISCUSSION

In this study, comprehensive capture probes were designed and used in conjunction with the capture enrichment method to sequence complete RSV and HuNoV genomes from clinical samples. These viruses represent two significant pathogens responsible for respiratory and gastrointestinal infections worldwide, requiring reliable methods for studying their genomic variability and evolution. The use of capture enrichment methodology overcomes any PCR primer design problems across the diverse viral strains and reduces non-target sequencing typically seen in standard RNA-seq.

Recently, Baier et al., designed their RSV capture probe set using a total of 1,101 complete genome sequences and used it to characterize the RSV-B outbreak in 2019 in four patients(16). Previously probe-based capture enrichment for HuNoV from human samples(26) and infected oysters (16) were reported. Brown et al.(21) reported the largest HuNoV probe set of the two studies which was designed using 622 norovirus partial or complete genomes and tested using different isolates of GI and GII(26). In this study, we report the custom-designed RSV probe set, based on 1,570 genomic sequences, covering 99.79% of targeted isolates, and the HuNoV probe set, designed from 1,376 sequences, covering 99.68% of targeted isolates which, to our knowledge, this represents the most comprehensive probe sets designed to date for sequencing the RSV and HuNoV.

Several process improvements such as sorting samples based on the Ct values (from high titer to low titer) on a plate during cDNA and library construction and arraying samples in alternate columns on a plate, were implemented to mitigate any potential contamination between samples. For target enrichment, to manage uneven sequence yields among samples, based on our previous experiences with SARS-CoV-2 enrichment, library pools were created based on Ct. values(27). While the uneven yields were still noted in these pools, enough reads were obtained for all 85 RSV and 47/55 HuNoV samples to generate full-length genomes.

A comparison between post-and pre-capture libraries for both RSV and HuNoV samples revealed that the percentage of reads aligning to the target virus genome (Table 1; Fig. 2), as well as the number of samples that resulted in full-length genomes (Fig. 3 and Fig. 4), was significantly higher in the post-capture libraries compared to the pre-capture libraries. Post-capture libraries showed 85.1% of reads mapping to the RSV genome, an 850x enrichment over the 0.08% in pre-capture libraries. In HuNoV samples, 40.8% of reads mapped post-capture, a 40.8x increase from the 1.15% in pre-capture libraries. These results are in line with previously reported probe-based enrichment methods for viral sequencing(27, 28).

Complete genomes were successfully assembled for all 85 RSV post-capture libraries, while only one complete genome was recovered from 24 pre-capture libraries. There were six samples under the ‘complete with low coverage’ genomes category and 17 samples with ‘incomplete’ genomes. (Table 1 and Table S1). Subtypes could be assigned to all 85 samples with 46 RSV-A subtypes and 39 RSV-B subtypes. RSVAB-WGS(29) is an amplicon-based protocol for RSV genome sequencing designed using 12 primers to cover both subtypes, producing PCR fragments of 1.5–2.5 kb. In 34 clinical samples, over 90% of the genome was recovered for Ct. values ≤ 25, while coverage dropped to 60–90% for Ct. 26-27 and 50% for Ct. above 27. In our study, we recovered full-length genomes from RSV A and B subtypes up to Ct. 30.

Complete genomes were successfully reconstructed for 47/55 HuNoV post-capture libraries. Among the remaining eight, two samples were categorized as ‘complete with low coverage’, four had ‘incomplete’ genomes and two samples failed to generate genome assemblies. These samples either had Ct higher than 33 (6/8 samples) or had failed in both post and pre-capture sequencing (2/8 samples), suggesting low viral titers or poor sample quality. As previous works have demonstrated, for reliable genome recovery the upper Ct threshold is approximately 30-33 cycles(27, 30). In the pre-capture set, only 18 out of 55 yielded complete genomes (Fig. 3), suggesting that capture enrichment is highly desirable.

The breadth of coverage at 20x depth was calculated to assess the efficiency of capture enrichment to enhance viral genome coverage depth (Fig 4). Notably, a substantial increase was observed in RSV, with both RSV-A and RSV-B samples exhibiting a dramatic post-capture rise in 20x coverage. HuNoV samples also displayed increased coverage post-capture, with remarkable coverage improvement across distinct genotypes, suggesting that the capture method offers significant benefits for RSV and HuNoV genome sequencing.

Both the results of this study and previous reports have shown that oligonucleotide capture methods show robust performance as the probes can tolerate variation in target sequences during enrichment, have overlapping designs, and can enrich from degraded samples, thereby greatly improving the chances of complete genome recovery(27, 28, 31).

The capture probes and the methodology described in this paper have been previously utilized to generate whole genome sequencing of both RSV and HuNoV clinical samples(32) https://www.biorxiv.org/content/10.1101/2023.05.30.542907v1.full.pdf). In the RSV study, 69 samples were collected longitudinally from HCT adults with normal (<14 days) and delayed (≥14 days) RSV clearance enrolled in a Ribavirin trial. Full-length genomes obtained from post-capture sequencing were analyzed across RSV-A or RSV-B to determine the inter-host and intra-host genetic variation and the effect on glycosylation(32).

In the HuNoV study, the evolutionary dynamics of human norovirus in healthy adults were studied using 156 HuNoV sequential samples from a controlled infection study(32) (https://www.biorxiv.org/content/10.1101/2023.05.30.542907v1.full.pdf).

Complete genomes were assembled for 123 of 156 samples (79%) including 45% of samples with Ct values below the limit of detection (>36 cycles) of the GI.1 genotype and collected up to 28 days post-infection. Non-synonymous amino acid changes were observed in all proteins, with capsid VP1 and nonstructural protein NS3 showing the highest variations. These findings indicate limited conserved immune pressure-driven evolution of the GI.1 virus in healthy adults and highlight the utility of capture-based sequencing to understand HuNoV biology.

Studying viral ORF expression is important to understanding viral pathogenesis, differentiation factors between subtypes, and the effects of genomic mutations on gene function including vaccine development. The RSV genome codes for 11 viral proteins, including three transmembrane glycoproteins G, F, and SH; matrix protein (M) and two transcription/replication regulating proteins (M2-1 and M2-2); three proteins related to nucleocapsid (N, P, L), and lastly two non-structural proteins NS1 and NS2(33). There are multiple reports of RSV ORF expression analysis where earlier studies suggested a gradient of gene transcription across the genome. ORF NS1 had the highest and ORF L had the lowest expression. Later reports demonstrated non-gradient mRNA levels, with the highest expression levels of the attachment ORF G(34–36). Differential patterns in RSV ORF expression in genotypes are also known(37). None of these studies used data from capture-enriched libraries that provide higher efficiency in RSV sequence recovery directly from patient samples.

Here we for the first time demonstrated the use of RSV sequence data generated from strand-specific libraries to study ORF expression. RSV is a negative-sense RNA virus and the ORFs are positive-strand mRNAs therefore, the reads from a strand-specific library derived from the sense strand (mRNA) will map onto the antisense strand of the reference genome, while those obtained from the genomic RNA map onto the sense strand.

While the ORF expression between post-capture and pre-capture libraries showed similar trends (Fig. 5) differences in ORF expression were not observed in a substantial number of both the RSV-A and RSV-B pre-capture libraries. This is not surprising given the low percentage of viral reads observed in these libraries. These results strongly suggest that the capture methodology significantly increased our ability to analyze ORF expression patterns without inducing any technical biases. Additionally, ORF expression differences were also noted between the two subtypes (Fig. 5). RSV-A subtype samples showed the highest expression in transmembrane ORFs SH and G, while ORFs M2-2 and M2-1 showed the lowest expression. RSV-B subtype samples had the highest expression in ORFs G and M and the lowest in ORFs M2-2 and N. Such genotype-specific differences were also reported by our group as well as others(17, 38). ORF gene expression generated from this approach can be utilized to investigate differences in viral gene expression *in vitro* within organoid models across various strains and hosts aiding in the study of RSV pathogenesis.

ORF analysis in HuNoV samples is not possible using the short reads generated in this study, as both the genome and ORFs in HuNoV are positive-strand RNA. Further, unlike the SARS-CoV-2 genome, where each ORF has a 5’ leader sequence, there are no such key ORF sequence differentiators in HuNoV that could be used to identify reads specifically originating from ORFs. Long-read sequencing data is recommended to identify and analyze HuNoV ORF expression profiles.

In conclusion, we describe two comprehensive probe sets and the capture enrichment methodology to successfully recover complete genomes from diverse genotypes of two important human viral pathogens. The methodology described to obtain the complete genome sequences is already in use to study viral genome evolution in these viruses. This type of sequencing data is also useful, as demonstrated here, in studying the RSV ORF expression patterns.

## MATERIALS AND METHODS

### Samples used in this study

RSV samples are part of active surveillance of pediatric acute respiratory illness (ARI) through the CDC’s New Vaccine Surveillance Network (NVSN) (https://www.cdc.gov/nvsn/php/about/index.html).

RSV-positive samples were collected from patients enrolled at the Houston NVSN site only. Mid-turbinate nasal and throat swab samples were obtained after informed consent was obtained verbally from the parent/guardian of the eligible children. Institutional review board approval was obtained locally from Baylor College of Medicine <u>(H-37691)</u> and at the CDC. HuNoV positive stool samples were collected as part of a controlled human infection model for GI.1 virus(39) as well as residual stool samples that were tested for gastrointestinal pathogens at Texas Children’s Hospital under an IRB-approved protocol.

In total 85 RSV samples and 55 HuNoV samples were characterized. All 85 RSV samples and 49/55 HuNoV are collected from patients while the remaining 6 HuNoV samples are from HuNoV-infected human intestinal organoids.

### RNA isolation

For the 85 RSV samples, approximately 200ul of each primary sample was extracted using the PureLink Pro Viral 96 DNA/RNA extraction kit (Thermo 12280096A) following the manufacturer’s instructions. Samples were eluted in 100ul.

For the 55 HuNoV stool samples, three RNA extraction kits were used starting with 0.2g of primary sample. For 33 samples, the MagAttract PowerMicrobiome DNA/RNA extraction kit (Qiagen 27500-4-EP) and for 16 samples, the AllPrep PowerFecal Pro DNA/RNA extraction kit (Qiagen 80254) was used. For the 6 HuNoV infected human intestinal enteroids, RNA was isolated using the MagMAX-96 viral RNA isolation kit. Samples were eluted in 100ul.

### Viral titer quantification

Real-time qPCR of RSV was performed using primers targeting the N gene as previously described(40).

HuNoV titers were assessed by reverse transcription-quantitative polymerase chain reaction (RT-qPCR), using the qScript XLT One-Step RT-qPCR ToughMix reagent with ROX reference dye (Quanta Biosciences). The primer pair and probe COG2R/QNIF2d/QNIFS(41) was used for GII genotype and NIFG1F/V1LCR/NIFG1P(42) was used for GI.1 genotype. Per sample Ct values can be found in Table S1.

### Capture probe design

The RSV probe set size was 23.77Mb and was designed based on 1,570 publicly available genomic sequences of RSV isolates. There are 87,025 unique probes of 80 bp length covering 99.79% of the targeted RSV isolates. The HuNoV probe set size was 9.6Mb and was designed based on 1,376 publicly available genomic sequences of HuNoV isolates, there are 39,300 unique probes of 80 bp length covering 99.68% of the targeted HuNoV isolates. The GenBank IDs for the references can be found in the capture design files of both RSV and HuNoV (see Table S3 and Table S4).

### cDNA preparation

Samples were processed in alternate columns on a 96-well plate and sorted from top left to bottom right from the highest titer to the lowest titer, because these libraries were prepared for capture enrichment, rRNA depletion, or Poly A+ RNA isolation steps were not performed.

### Capture enrichment and sequencing

RSV and HuNoV cDNA were hybridized in separate pools with biotin-labeled RSV and HuNoV capture probes. The 85 RSV samples were enriched in three library pools consisting of 24 samples with Ct. values 17 to 21.5, 31 samples with Ct. values 21.8 to 25, and 30 samples with Ct. values 25.1 to 29.9 along with samples with Ct ND. The 55 HuNoV libraries were grouped as three pools, with one pool containing 14 samples with Ct. values between 21.5 - 25.7 and a second pool containing 13 samples with Ct. values between 26.3 and 34.5 along with samples with Ct values ND, and the third pool containing 28 samples with Ct. values between 20.2-34.88.

All six pools of cDNA libraries were incubated at 70°C for 16 hours followed by enrichment PCR as previously reported(27). The amount of each cDNA library pooled for hybridization and post-capture amplification of 12-20 PCR cycles was determined empirically according to the virus Ct values. Between 1.8–4.0 μg pre-capture library was used for hybridization with the viral probes and the post-capture libraries were sequenced on Illumina NovaSeq S4 flow cell, to generate 2×150 bp paired-end reads. Pre-capture libraries for 24 RSV samples and all 55 of the HuNoV samples were also sequenced.

### RSV and HuNoV genome assembly

Following sequencing, raw data files in binary base call (BCL) format were converted into FASTQs and demultiplexed based on the dual-index barcodes using the Illumina ‘bcl2fastq’ software. Demultiplexed raw fastq sequences were processed using BBDuk (https://sourceforge.net/projects/bbmap/) to quality trim, remove Illumina adapters, and filter PhiX reads. Trimmed FASTQs were mapped to a combined PhiX (and human reference genome database (hg38) using BBMap (https://sourceforge.net/projects/bbmap/) to determine and remove human/PhiX reads. Trimmed and host-filtered reads were processed through VirMAP (24) to assemble complete RSV or HuNoV genomes. The VirMAP summary statistics include information on reconstructed genome length, the number of reads mapped to the reconstruction, and the average coverage across the genome.

HuNoV genome reconstructions were genotyped via the CDC-developed Human Calicivirus Typing Tool (https://calicivirustypingtool.cdc.gov/bctyping.html). Final reconstructions were manually inspected using Geneious Prime® 2022.1.1 and aligned against the relevant HuNoV or RSV reference genomes to determine the quality of assemblies. The breadth of coverage at 20x depth was calculated by re-aligning the raw reads reference genome (RSV) or individual sample-assembled genome (HuNoV) using BWA MEM https://arxiv.org/abs/1303.3997 (version 0.7.17- r1188) with standard parameters. Coverage for each sample was assessed using “samtools depth” (version 1.6), applying a mapping quality filter of 20 phred scores (-q 20). Downstream analysis of summary statistics was done using R (https://www.r-project.org/).

### RSV expression profile analysis

VirMAP(43) trimmed reads from both the pre-and post-capture datasets were mapped to RSV-A ON and RSV-B BA reference genomes(32), according to the sample genotypes, using BBMap version 39.01. Gene annotation for the reference genomes ON and BA was conducted using VIGOR(44). Since RSV is a negative-stranded RNA virus, read pairs with read 1 mapped to the negative strand are from the viral genome, while read pairs with read 1 mapped to the positive strand of the reference genome are from the viral mRNAs. Read pairs were assigned to each gene using featureCounts version 2.0.1(45) with “-s 1 -p” options for counting read pairs mapped to the positive strand of the reference genome. The read pair counts assigned to each gene were then normalized to the number of read pairs per kb gene length and per million mapped reads (FPKM) and plotted using the R ggplot2 package (https://ggplot2.tidyverse.org/).

## ACKNOWLEDGEMENTS

The authors are grateful to the production teams at The Alkek Center for Metagenomics and Microbiome Research and Human Genome Sequencing Center for data generation. We would also like to thank Frederick Neill from the HuNoV group. This work was supported by the National Institute of Allergy and Infectious Diseases (Grant#1U19AI144297). No additional external funding was received for this study.

## AUTHOR CONTRIBUTIONS

S.J.C and H.D: Conceptualization

S.V.B, A.S, D.P.A, S.J.C and H.D: Writing and Analysis.

S.V.B, D.P.A, H.C, K.K, Z.M, G.W, N.E, F.M, J.H, V.K.M, Q.M: Data Generation

A.S: Project Administration

Q.X, A.S, K.W, R.S, D.H, F.S, Z.K, G.A.M, V.A, P.O, S.R, R.A, M.E, D.M.M: Review & Editing

R.A.G, J.F.P: Funding Acquisition

## DATA AVAILABILITY

Complete genomes and raw fastq files for the samples used in this study are being uploaded to NCBI GenBank and SRA, respectively, under BioProjectID XXXX. Analysis and figure code is available at the following GitHub link: https://github.com/BCM-GCID/Capture_benchmarking_paper

